# A latitudinal gradient of reference genomes

**DOI:** 10.1101/2024.07.09.602657

**Authors:** Ethan B. Linck, Carlos Daniel Cadena

## Abstract

Global inequality rooted in legacies of colonialism and uneven development can lead to systematic biases in scientific knowledge. In ecology and evolutionary biology, findings, funding and research effort are disproportionately concentrated at high latitudes while biological diversity is concentrated at low latitudes. This discrepancy may have a particular influence in fields like phylogeography, molecular ecology and conservation genetics, where the rise of genomics has increased the cost and technical expertise required to apply state-of-the-art methods. Here we ask whether a fundamental biogeographic pattern—the latitudinal gradient of species richness in tetrapods—is reflected in available reference genomes, an important data resource for various applications of molecular tools for biodiversity research and conservation. We also ask whether sequencing approaches differ between the Global South and Global North, reviewing the last five years of conservation genetics research in four leading journals. We find that extant reference genomes are scarce relative to species richness at low latitudes, and that reduced-representation and whole-genome sequencing are disproportionately applied to taxa in the Global North. We conclude with recommendations to close this gap and improve international collaborations in biodiversity genomics.

## Introduction

Scientific knowledge reflects both reality and the economic and social conditions that shape the practice of research (Hull 1990; Logino 2002; Stephan 2012). The coincidence of the Scientific Revolution and imperial expansion in Western Europe in the 16th through 18th centuries led to the accumulation and concentration of global capital in institutions of higher learning in the United Kingdom, France, Spain, the Netherlands, and Germany, funding foundational investigations in the natural sciences. Industrialization and the rise of capitalism in the 19th century stoked the boiler of the research endeavor, permitting the continued expansion of universities and birthing a learned aristocratic class with disposable time and income to commit to natural history (Opitz 2004; Opitz 2006). This legacy persists to this day: Research expenditures concentrated in a handful of majority Anglophone countries (May 1997), which in turn produce a disproportionate amount of scholarship (May 1997; King 2004) and often drive citation networks (Pasterkamp et al. 2007; Meneghini et al. 2008).

Ecology and evolutionary biology (EEB) face a unique challenge in light of global scientific inequality, as while funding and research effort are disproportionately concentrated at high latitudes (Melles et al. 2019), biological diversity is concentrated at low latitudes (Willig et al. 2003). The consequences of this discrepancy are many, including systematic biases in research effort (Titley et al. 2017) and taxonomy (Freeman & Pennell 2021), major gaps in our understanding of natural history and species distributions (Collen et al. 2008; Feely & Sillman 2011), and a blinkered view of patterns of diversity and diversification across the tree of life (Reddy 2014; Cornwell et al. 2019). Increased research attention on tropical ecosystems has therefore been a stated priority for decades (Collen et al. 2008; Nori et al. 2020), but despite some promising strides towards international collaborations (Perez et al. 2018), partners in the Global North frequently set research agendas (Bradeley 2008; Asase et al. 2022), leading to accusations of parachute science and persistent gaps in data quantity, quality, and type (Stocks et al. 2008; Soares et al. 2023).

EEB disciplines where data collection and analysis are equipment- and cost-intensive are especially likely to show increased discrepancies between the Global North and Global South. In the 1960s and 70s, assays of molecular genetic markers in non-model organisms laid the groundwork for phylogenetic systematics to shift away from a traditional focus on morphology, increasing statistical power and requiring increasingly expensive tools and specialized skills (Hillis et al. 1996). Concurrently, the rise of phylogeography brought an evolutionary perspective to population biology, altering views on the fundamental units of management and conservation (Avise 2000). In the late 2000s and early 2010s, reduced representation high-throughput sequencing using library preparation methods such as RADseq and target-capture began to proliferate as an alternative to traditional Sanger sequencing (Eklbom et al. 2011; Lemmon & Lemmon 2013), boasting the advantages of rapidly generating much larger datasets and requiring little to no prior information about a focal taxon’s genome.

A third revolution is currently underway, as low-coverage whole-genome resequencing (hereafter WGS) begins to supplant reduced representation methods in some fields and taxa (Ellegren et al. 2014; Toews et al. 2016). Unlike RADseq, target capture, and related approaches, WGS typically requires a high-coverage reference genome with which to align samples. Because sequencing an individual at sufficient depth remains costly—and because assembling such large data is computationally intensive—generating high-quality novel reference genomes remains difficult for many labs and, by extension, species.

The problem may be particularly acute in conservation biology where decades of concern about the so-called “conservation genetics gap” have highlighted a persistent disconnect between the importance managers place on genetic information and its actual use in decision making (Taylor et al. 2017). Hypotheses to explain the conservation genetics gap include skepticism about its importance, the specialized knowledge required for analysis and interpretation, and cost (Hoban et al. 2013), though some recent discussion suggests the term itself obfuscates by lumping the diverse set of obstacles (or “spaces”) between research and implementation (i.e., Toomey et al. 2017; see solutions reviewed in Hogg et al. 2024). At low-resource institutions in the tropics a broader shift towards WGS approaches may further discourage the development of the field in the very regions where the greatest number of critically endangered species are found (Vamosi & Vamosi 2008; Bertola et al. 2024), especially if technological advances become publication requirements at high impact and broadly read journals.

Here we ask whether a fundamental biogeographic pattern—the latitudinal gradient of species richness in tetrapods—is reflected in reference genomes available through the US National Institute of Health’s National Center for Biotechnology Information Genome Browser, an important data resource for various applications of molecular tools for biodiversity research and conservation. We hypothesized that NCBI contributor affiliations combined with inequities in economic development and access to scientific resources would lead the number of species with assembled genomes to be greatest in the temperate zone, not the tropics. We also ask whether sequencing approaches differ between the Global South and Global North, reviewing the last five years of conservation genetics research in four leading English-language journals. We conclude by discussing strategies to improve international collaborations in biodiversity genomics and boost the representation of species from the Global South in sequence databases.

## Methods

### Reference genomes, species richness, and latitude

We used the National Center for Biotechnology Information (NCBI) Datasets command-line tools v.16.19.0 (O’Leary et al. 2024) to download taxonomy metadata for the subset of species with an assembled reference genome in the following taxa: birds (Class: Aves), mammals (Class: Mammalia), squamates (Order: Squamata), amphibians (Class: Amphibia), turtles (Order: Testudines), crocodilians (Order: Crocodilia) and tuataras (Order: Rhynchocephalia). We selected these groups—together comprising extant tetrapods—to provide a snapshot of animal diversity in relatively well-studied clades with different ecologies and evolutionary histories, while restricting the total dataset to a computationally manageable size. From this initial list we retained species with an exact match to the Global Biodiversity Information Facility’s (GBIF) Backbone Taxonomy using rgbif v.3.8.0 (Chamberlain et al. 2024) and downloaded all observations of each backed by georeferenced voucher specimens in natural history museum collections (NHCs), excluding those without coordinates and those flagged for geospatial issues (n= 3,006,946). We repeated this process for all species in each higher-level taxon represented in our list of reference genomes (i.e., downloaded metadata for all georeferenced tetrapod specimens on GBIF; n= 9,303,258). DOIs for each download are available in the References section below (GBIF.org 2024a; GBIF.org 2024b; GBIF.org 2024c; GBIF.org 2024d; GBIF.org 2024e; GBIF.org 2024f; GBIF.org 2024g; GBIF.org 2024h).

Filtering these aggregated datasets to contain only species with 10 or more specimen records, we generated convex hull polygons for each as a coarse approximation of their geographic distribution using the R package sf v.1.0-16 (Pebesma 2018; Pebesma & Bivand 2023). Overlaying these on a shapefile of Earth’s landmasses from rnaturalearth v.1.0.1 (Massicotte & South 2024), we calculated species richness as the number of overlapping convex hulls in 2-degree x 2-degree grid cells, statistically standardizing this value by subtracting observed mean global species richness and dividing by its standard deviation. We subtracted the number of species with reference genomes from total species richness to determine the regions with the largest representation gap in genomic resources, again standardizing the difference. To assess the significance and slope of a correlation between species richness and the absolute value (or modulus) of latitude in decimal degrees, we performed simple linear regressions in R v.4.4.0 (R Core Team 2024), analyzing species with reference genomes and our full dataset separately.

### Literature review

To evaluate how the geography of authorship might impact sequencing strategy of studies in conservation biology, we performed a restricted Web of Science literature search on 29 June 2024 for English-language conservation genetics papers published in the last five years in the journals *Conservation Genetics*, *Molecular Ecology*, *Journal of Heredity*, and *Conservation Biology,* selected for frequently publishing empirical work on non-model organisms. We used the queries ‘SO=”Conservation Genetics”’ and ‘SO=(“Molecular Ecology” OR “Journal of Heredity” OR “Conservation Biology”) AND (TS=“Conservation Genet*” OR KP=“Conservation Genet*” OR TI=“Conservation Genet*”’), excluding reviews, genome announcements, meta-analyses, preprints, and studies that were purely simulations. Our criteria aimed to achieve a tractable sample size for careful study (<1000 papers) while covering the period in which WGS became commonly used for the conservation genetics of non-model organisms (Fuentes-Pardo & Ruzzante 2017; Hohenlohe et al. 2021).

We then manually reviewed each study, first assigning the home institution of its first and last author to the Global North, Global South or both (i.e., joint affiliations) using the 2024 UN Trade and Development Classifications. Because the number of middle authors varied widely across our sample, we assessed their affiliations on a binary basis, indicating only whether a contributor from an institution from the Global South was present outside of the lead and senior positions. Synthesizing these data, we assigned papers to mutually exclusive groups based on whether they included one or more Global South authors or only Global North authors. Next, we categorized each study’s sequencing approach as reduced representation, WGS, Sanger sequencing, microsatellites, or other, and described its overall focus using tiered categories based on discussion in Bertola et al. 2024. These tiers were: 1) Taxonomy / systematics, identification, or sexing; 2) Phylogeography / population genetic structure, estimating genetic diversity, and inferring demographic history; and 3) Detecting outlier loci, quantifying runs of homozygosity, and evaluating adaptive potential. When studies employed more than one sequencing approach or addressed goals belonging to multiple tiers, we assigned them to a single category based on their most data-intensive method or question. To explore geographic patterns in sequencing effort, we assessed whether each study’s taxonomic sampling included 1) at least one species distributed in the Global South and 2) at least one species distributed in the Global North. Because some studies included multiple taxa and some species are broadly distributed or migrate between regions, these categories were not mutually exclusive. To evaluate whether geographic representation in conservation genetics changed over the period covered by our review, we performed logistic regression using the stats package R v.4.4.0, treating the presence or absence of an author from the Global South as a binary outcome variable and year as the sole independent variable.

## Results

### Sampled Gradients of Species Richness

Our list of tetrapod reference genomes from NCBI included 1159 bird species, 795 mammal species, 123 amphibian species, 39 turtle species, 6 crocodile species, and the tuatara (*Sphenodon punctuatus*). This total represented 6.8% of the 30,832 tetrapod species with a georeferenced preserved specimen record on GBIF. Species with a reference genome were associated with 3,048,136 specimens, or 40.7% of the 7,478,867 tetrapod specimens with available metadata. Following filtering out species with 10 or fewer observations, we retained 1,859 species with a reference genome, or 8.6% of the 21,583 tetrapod species meeting the same criteria.

Minimum tetrapod species richness in both datasets was 1. Maximum species richness calculated from the reference genome dataset was 705, occurring in a grid cell centered on south Florida, USA (**Figure 1C**). Maximum species richness in the full dataset was 2698, occurring in a grid cell centered on the western Amazon basin and east slope of the Andes in Ecuador near the Peruvian border (**Figure 1C**); this was also where the greatest gap between sequenced species and total species richness was observed (**Figure 1A**).

**Figure 1.**
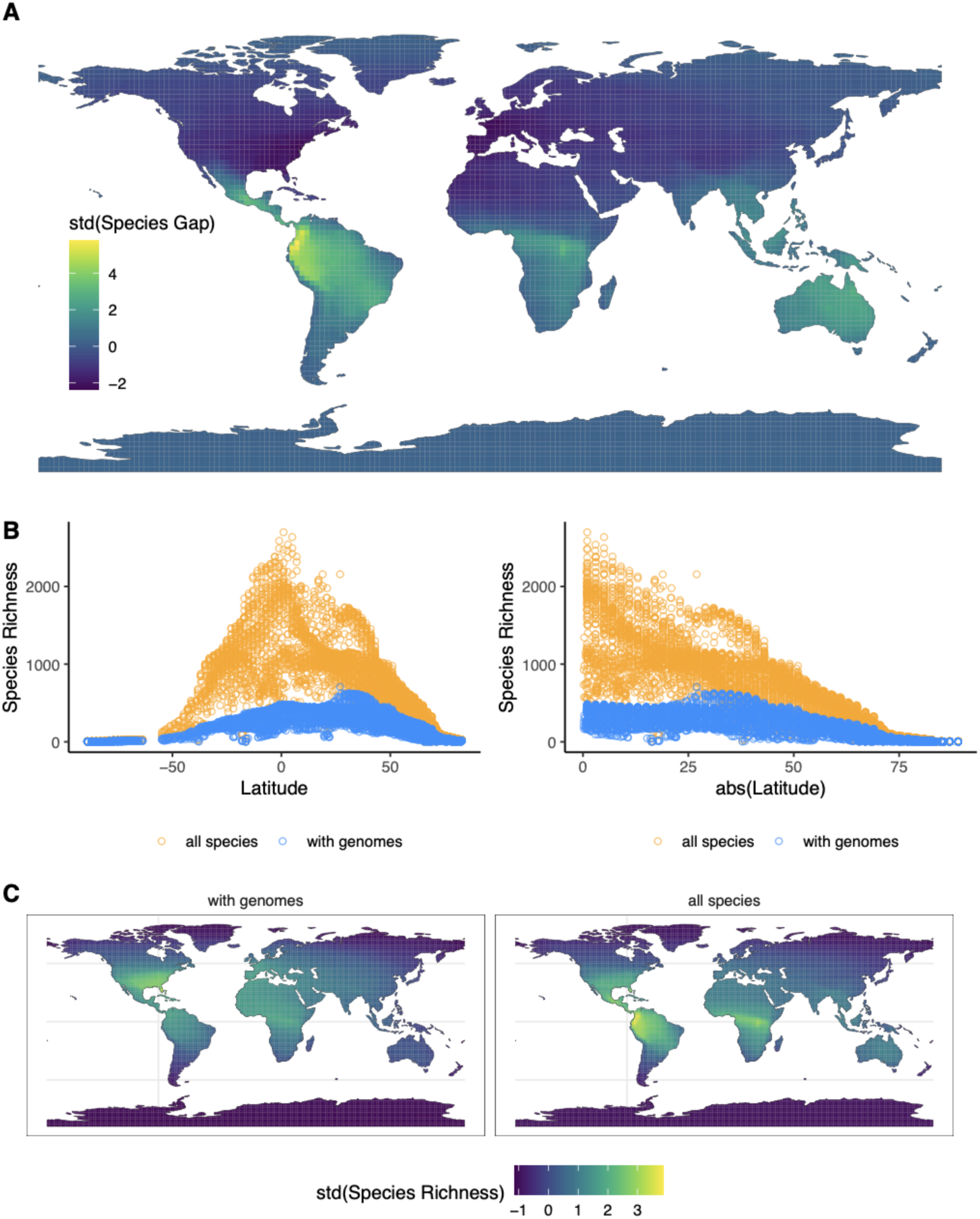
A) Tropical species are underrepresented among available reference genomes. Colors reflect the standardized difference between total species richness and the number of species with an assembled reference genome on the NCBI genome browser. Negative values indicate grid cells falling below calculated global mean species richness. B) Richness among species with reference genomes does not reflect global patterns of biodiversity. Blue circles represent total local species richness in 2-degree by 2-degree grid cells, while gold circles represent richness of species with assembled reference genomes. C) Global patterns of species richness calculated from species with reference genomes and all species with NHC specimen records on GBIF, respectively.

Species richness was negatively related to the absolute value latitude in both regressions, albeit with a much steeper slope when data from all tetrapod species were included (reference genomes only, β = −4.892, adjusted *R*^2^=0.6439; p<0.001; full data, β = −18.65, adjusted *R*^2^=0.7856 p<0.001) (**Figure 1B**). Because data visualization indicated there might be a distinct breakpoint in the relationship at mid latitudes—potentially reflecting a transition from the influence of sampling effort effects to true biogeographic signal—we fit an additional piecewise linear regression model with the R package segmented v.2.1-0. This model identified a breakpoint at 39.819, fitting a segment with a slope of β = −0.312 before it, and a segment with a slope of β = −7.3634 after it (p=0.0408; adjusted *R*^2^=0.7184).

### Literature Review

After excluding papers that did not meet our stated criteria, we reviewed 394 empirical conservation genetics articles published between 1 January 2020 and 29 June 2024. This list included 345 papers from the journal *Conservation Genetics,* 11 papers from *Journal of Heredity*, 35 articles from *Molecular Ecology,* and 3 articles from *Conservation Biology*. Of these, 62 included a first or senior author from the Global South; 40 more included Global South scientists as middle authors. Nearly 87% (n=342) of papers had a first or senior author from the Global North. Ninety-eight included sampling from a focal taxon or focal taxa in the Global South, with 277 sampling a taxon or taxa from the Global North. From 2020 to 2024, the odds of a paper including an author from the Global South did not significantly increase (log-odds 1.1096; 95% CI −0.5864-0.2667; p=0.210). Microsatellites were the most used sequencing strategy among Global South authors and in studies of Global South taxa, while reduced representation genome sequencing approaches were most common in the Global North. For authors and taxa in both the Global South and the Global North, Tier 2 studies (phylogeography / population genetic structure, estimating genetic diversity, or inferring demographic history) were most common. Further details are provided in **Table 1** and **Table 2**.

**Table 1.** Summary of regional authorship affiliation, sequencing strategy, and sampled focal species range for empirical conservation genetics papers from 2019-2024 in four leading journals. Integers indicate the total number of studies in each category, while numbers in parentheses refer to its proportion out of all reviewed articles (n=394). Regional authorship affiliation is multually exclusive, while the distribution of focal taxa is not, as studies could involve multiple taxa, broadly distributed taxa, or long-distance migrants. Papers were assigned to a sequencing strategy based on the most data-intensive approach they employed (i.e., a study applying both Sanger sequencing and microsatellites would be assigned to the ‘Microsatellites’ category.)

| **Sequencing Strategy** | **Global South Author** | **Only Global North Authors** | **Global South Taxon** | **Global North Taxon** |
| --- | --- | --- | --- | --- |
| Sanger | 22 (0.0558) | 25 (0.0634) | 26 (0.0660) | 26 (0.0659) |
| Microsatellites | 43 (0.1066) | 105 (0.2690) | 45 (0.1142) | 108 (0.2741) |
| Reduced Representation | 29 (0.0736) | 129 (0.3299) | 31 (0.0787) | 131 (0.3325) |
| WGS | 5 (0.0013) | 24 (0.0609) | 11 (0.0152) | 18 (0.0279) |
| Other | 4 (0.0101) | 8 (0.0203) | 4 (0.0102) | 8 (0.0203) |

**Table 2.** Summary of regional authorship affiliation, study goals, and sampled focal species range among reviewed papers. Study goals refer to broad tiers of research questions with increasing data requirements. Papers were assigned to each on the basis of the most data-intensive analysis they employed (i.e., a paper inferring population genetic structure and identifying loci under selection would be assigned to the tier 3). As in Table 1, ‘Global South Author’ and ‘Only Global North Authors’ are mutually exclusive categories, while ‘Global South Taxon’ and ‘Global North Taxon’ are not.

| **Study Goals** | **Global South**<br>**Author** | **Only Global**<br>**North Authors** | **Global South**<br>**Taxon** | **Global North**<br>**Taxon** |
| --- | --- | --- | --- | --- |
| 1. Taxonomy / systematics, identification, or sexing | 9 (0.0228) | 19 (0.0635) | 11 (0.0279) | 17 (0.04315) |
| 2. Phylogeography / population genetic structure, estimating genetic diversity, and inferring demographic history | 90 (0.2284) | 241 (0.6192) | 102 (0.2589) | 248 (0.6294) |
| 3. Detecting outlier loci, quantifying runs of homozygosity, and evaluating adaptive potential | 4 (0.0106) | 31 (0.0786) | 4 (0.0102) | 31 (0.0787) |

## Discussion

Reference genomes available through the US National Institute of Health’s National Center for Biotechnology Genome Browser fail to reflect the overwhelming concentration of tetrapod species richness in the tropics and are strongly biased towards species at mid-latitudes in the Northern Hemisphere (**Figure 1**). This pattern is almost certainly a result of global inequalities in economic development and its resulting effects on research productivity (May 1997; King 2004). Its consequences will likely include increasing an already profound methodological gap in sequencing approaches between molecular ecologists in the Global North and the Global South (**Table 1**).

Our analysis contrasts patterns inferred from both a traditional source of biodiversity data—vouchered specimens in NHCs—and a contemporary genomic resource archive, the National Institute of Health’s NCBI Genome Browser. Our description of the latitudinal gradient of total tetrapod species richness is broadly similar to other recent macroecological studies of the phenomenon (Roll et al. 2017; Quintero et al. 2023), with global hotspots concentrated in northwest South America and Central Africa (**Figure 1C**) and an approximately monotonic decline in latitudinal species richness maxima from the equator to the poles. In contrast, the latitudinal gradient of richness of tetrapod species with reference genomes is flattened, showing only moderate declines at high latitudes and a mid-latitude peak in species richness in the Northern Hemisphere (**Figure 1B**).

This difference is especially notable because we made no effort to correct for disparities in historical specimen collection across latitude, with the consequence that our ‘true’ species richness gradient significantly underestimates biodiversity in the tropics. Across longitude, our analysis appears to underestimate diversity in East Asia, Indonesia, and Oceania (Quintero et al. 2023), likely due to both the coarse grain of our study and the region’s greater distance from the large NHCs in Europe and North America that are the backbone of curated GBIF data. Geopolitical factors complicate a simple interpretation of these results: use of GBIF and NCBI by scientists outside the Anglosphere may be limited by data sovereignty laws or other impacts of economic and military rivalries. Regardless, species with publicly available reference genomes as of July 2024 are more reflective of socioeconomic conditions than biogeographic reality.

If both natural history collections and contemporary bioinformatics resources reflect historical inequalities in development and scientific capacity, why do data from the former better approximate the latitudinal gradient of species richness? Part of the answer lies in their different goals: while the mission of many NHCs is to explicitly catalog and archive regional or global biodiversity, the NCBI Genome Browser is typically used as a repository for open data publication requirements and is less often an end unto itself. Another possibility is that researchers looking to obtain genetic material from the tropics face unique obstacles to their work. As the birth and golden age of NHCs coincided with the heyday of Western colonialism (De Vos 2007; Quintero Toro 2012), scientific collecting was less restricted by concerns of either sovereignty or Indigenous land tenure, let alone local or international regulations on the transfer of biological samples (Kahn & Tyagi 2021; Sherkow et al 2022). Relatedly, one early reader of this manuscript pointed to the added challenge of obtaining high-quality tissue samples in hot, humid conditions.

We do not believe access to sources of DNA is a major obstacle: Science is rich with examples of researchers overcoming far greater hurdles to their work, and extant frozen tissue collections at institutions in the both the Global South and Global North include thousands of samples awaiting sequencing. Instead, we suggest that evolving norms and persistent obstacles of cost have led many relatively well-resourced scientists in the Global North to prioritize generating reference genomes for local taxa. In spite of rapid declines in the per base pair cost of whole-genome sequencing (Lou et al. 2021), high-coverage sequencing remains a significant expense: while averages are hard to come by in this increasingly privatized sector, one of us recently paid ∼$13,000 USD (September 2023) for high-depth long-read and HiC sequencing of a North American passerine bird from an industry contractor. Even in North America, this figure likely pushes small, single PI labs to prioritize investing in generating resources for species at the center of their research program, or otherwise likely to provide long-term utility—which are often those in their own backyards.

In the spirit of the traditional mission of NHCs, the past decade has seen several interrelated, international initiatives to increase taxonomic diversity in high-quality reference genomes (e.g. O’Brien et al. 2014; Koepfli et al. 2015; Cheng et al. 2018; Rhie et al. 2021). Of these initiatives, the Earth BioGenome Project (EBP)—a “moonshot” endeavor to sequence all Eukarya within ten years—is by design the most ambitious, bringing its smaller-scale predecessors together under a common set of goals and standards (Lewin et al. 2018). These collaborations have had a profound impact on biodiversity genomics: As of March 2021, EBP estimated at least one species from 15.6% of all Eukaryotic families had been sequenced, marking notable progress towards its Phase I goal of complete family-level representation within 3 years. Though this focus on phylogenetic diversity has introduced its own bias by undersampling lineages with high diversification rates, the initiative has nonetheless helped to close what was surely an even larger gap in empirical patterns of species richness and species richness represented by NCBI. In some cases, EBP or the initiatives under its umbrella have also provided researchers with early access to draft assemblies of nonmodel organisms (e.g., Linck et al. 2020) or with support and resources to produce new assemblies (e.g., Cadena et al. 2024), a scenario that suggests the resource gap may be slightly less dire in practice than reported here.

Yet in a world of limited time, finite resources, and incompletely described biodiversity, sequencing at scale is not immune to its own biases. For example, a recent publication introducing an attempt to generate reference genomes for all vertebrates (Rhie et al. 2021) included 127 authors affiliated with 102 institutions. Of these, only 13 are in the Global South: 4 in China, 3 in Korea, 2 in Malaysia, 1 in Singapore, 1 in Qatar, and 1 in Colombia. No authors or institutional affiliations from Africa, Indonesia, or Oceania outside of Australia and New Zealand were included. To date, institutions in the Global North have generated most animal (Hotaling et al. 2021) and plant (Marks et al. 2021) genomes available on NCBI’s GenBank, a situation mirrored by an analysis of squamates alone by Pinto et al. (2023). Though understandable considering the current distribution of scientists and resources—though not of species—the imbalance seems likely to perpetuate representational biases in the near-term.

Current efforts by international collaborations like the Amphibian Genomics Consortium to increase geographic representation and to offer support and opportunities to researchers from developing countries and underrepresented groups (Kosch et al. 2024) are steps in the right direction in the path to make the field of biodiversity genomics more equitable. We are particularly enthusiastic about EBP’s model of pairing targeted sequencing efforts with capacity building workshops (see Sharaf et al. 2024 for a recent example from 11 African countries). In line with the Convention on Biological Diversity’s Nagoya Protocol (Secretariat of the Convention on Biodiversity 2011), another critical dimension of the conversation about equitable generation of encyclopedias of reference genomes is the need for researchers to build strong partnerships with Indigenous peoples and other local communities, allowing them to participate in and benefit from the different phases and products of sequencing projects (Ambler et al. 2020; Colella et al. 2023; Mc Cartney et al. 2023).

If tropical species are underrepresented on NCBI, we would expect that they are only rarely studied using whole-genome resequencing (and other sequencing strategies dependent on a reference genome). Our review of conservation genetics papers published in *Molecular Ecology, Journal of Heredity, Conservation Genetics,* and *Conservation Biology* over the last five years suggests this is indeed the case. While the Global South / Global North binary and measures of human development more generally are only imperfectly correlated with latitude, their association nonetheless indicates that WGS is only rarely applied in conservation genetics studies in the tropics (n=6), and almost never by a leading researcher with a primary affiliation to a research institution in the region (n=1) (**Table 1**).

Similarly, for Global South scientists and for Global South focal taxa, microsatellites remain the most common molecular approach; for scientists and focal taxa in the Global North, reduced representation approaches dominate. We believe this is reflective of the expense and limited availability of high-throughput sequencing in the Global South, regardless of whether reads are assembled *de novo* or aligned to a reference. Lastly, we point out that across all categories, the vast majority of scientists (87%) and species (77%) in our sample originate in the Global North. Though our choice of North American or European-based journals precludes generalization, the primacy of the English language in global science (Montgomery 2013) leads us to interpret this as indicating the field of conservation genetics—let alone conservation genomics—is in its infancy in the tropics. We thus doubt the patterns described here would change appreciably if we were to include publications in other languages. Setting appropriate goals, targets, and indicators to effectively conserve and monitor global genetic diversity will require this situation to be remedied (Hoban et al. 2021).

We highlight discrepancies between available reference genomes and global biogeographic patterns to encourage increased, equitable collaboration between scientists in the Global North and Global South (see Sawchuk et al. 2024 for an African-based perspective on equity and inclusion in population genetics and ancient DNA studies). In light of this, we make three simple recommendations (see also Bertola et al. 2023). First, we encourage scientists from resource-rich institutions to consider allocating effort and funds towards generating reference genomes that serve the needs of managers and researchers in the Global South. Second, we support the continued development of multinational sequencing projects, but ask funders and senior personnel to increasingly consider prioritization, inclusion, and capacity building in areas of the world with rich biodiversity and limited resources to study it using genomic tools Third, we ask journals to consider issues of access and cost in editorial guidelines and decisions: While high-throughput sequencing is increasingly expected by editorial boards and reviewers at high-impact journals, it is not essential to address a variety of research questions (Bertola et al. 2023) and remains out of reach for most scientists residing where most of the world’s species occur. If ecology, evolution, and conservation aim to accurately catalog and effectively protect life on Earth, remedying inequalities in genomic resources should be a major priority.

## Data Accessibility

Processed GBIF datasets, NCBI metadata, and a digital notebook containing code to perform these analyses and generate Figure 1 are available at https://github.com/elinck/lat_grad_genome and from Data Dryad (https://doi.org/10.5061/dryad.2v6wwpzxh).

## Acknowledgments

We thank Marty Kardos for the invitation to participate in this special issue. Carolyn Hogg and an anonymous reviewer provided helpful feedback on the study. E.B.L. expresses appreciation for his mobile phone’s Wi-Fi hotspot, which was essential for submitting the initial draft of this manuscript from the field. Ongoing genomic work by C.D.C. is supported by the Facultad de Ciencias at Universidad de Los Andes through a *Programa de Investigación* (Convocatoria 2024-2026).

